# Evaluation of microRNA variant maturation prior to genome edition

**DOI:** 10.1101/2022.12.16.520740

**Authors:** Isabelle Busseau, Sophie Mockly, Élisabeth Houbron, Hedi Somaï, Hervé Seitz

**Author notes:** Corresponding authors; (I.B.) and (H.S.). Institut de Recherches Cliniques de Montréal (IRCM), Montréal, QC, Canada.

## Abstract

Assessment of the functionality of individual microRNA/target sites is a crucial issue. Genome editing techniques should theoretically permit a fine functional exploration of such interactions, allowing the mutation of microRNAs or individual binding sites in a complete *in vivo* setting, therefore abrogating or restoring individual interactions on demand. A major limitation to this experimental strategy is the influence of microRNA sequence on its accumulation level, which introduces a confounding effect when assessing phenotypic rescue by compensatorily mutated microRNA and target site. Here we describe a simple assay to identify microRNA variants most likely to accumulate at wild-type levels even though their sequence has been mutated. In this assay, quantification of a reporter construct in cultured cells predicts the efficiency of an early biogenesis step, the Drosha-dependent cleavage of microRNA precursors, which appears to be a major determinant of microRNA accumulation in our variant collection. This system allowed the generation of a mutant *Drosophila* strain expressing a *bantam* microRNA variant at wild-type levels.

## INTRODUCTION

microRNAs (miRNAs) are small regulatory RNAs which guide a repressive ribonucleoproteic complex onto specific RNA targets. miRNA targets are recognized by sequence complementarity, with most known targets in animals exhibiting a perfect complementarity to the miRNA “seed” sequence (nt 2–7), and imperfect complementarity to the rest of the miRNA sequence [1]. While computational tools, and high-throughput molecular biology methods, typically predict hundreds of RNA targets for each miRNA, *in vivo* genetics usually identifies just a few key targets responsible for miRNA-controlled phenotypes [2, 3, 4, 5, 6].

Such discrepancy between molecular biology and bioinformatics on one hand, and *in vivo* genetics on the other hand, is a central question in miRNA biology. The development of CRISPR/Cas9-mediated genome editing now offers a great opportunity to assess it rigorously, with the possibility to mutate miRNA genes, individual miRNA binding sites on target RNAs, or both of them simultaneously (offering the possibility to restore a single interaction with compensatory mutations in the miRNA and its target) [3]. It can be expected that such precise genetics experiments will disprove the biological functionality of many published miRNA/target interactions, as exemplified by the *bantam*/*enabled* interaction in *Drosophila* [7, 8].

*miRNA genes are typically transcribed as long precursors called “pri-miRNAs”. In animals (see* [1] for a review), most pri-miRNAs are cleaved in the nucleus by the Drosha endonuclease, liberating a short (*≈* 70 nt) stem-loop-folded “pre-miRNA”. The pre-miRNA is exported to the cytoplasm, where most pre-miRNAs are cleaved by the Dicer endonuclease: the cleavage reaction liberates the loop (which is quickly degraded) from the stem. The stem is composed of two small RNAs which are essentially complementary to each other. That duplex is then loaded on a protein of the Ago subfamily of the Argonaute family, and one strand is ejected and quickly degraded (see [9] for a review). In general for a given miRNA gene, the same RNA strand tends to remain associated to Ago (that strand is called the “miRNA”), while the other strand tends to be most frequently ejected and degraded (that other strand is called the “miRNA*”).

Known determinants of miRNA maturation from their pri-miRNA and pre-miRNA precursors rely on sequence and secondary structure motifs, none of them being located in the miRNA/miRNA* duplex [10, 11, 12]. But in addition to these well-defined motifs, miRNA maturation relies on the stability of the stem-loop structure, and mutations affecting the double-stranded structure of the stem may also perturb miRNA biogenesis. This dependency creates a major obstacle to the generation of useful genome-edited miRNA genes, complicating the interpretation of biological phenotypes when a mutated miRNA is not only modified in its sequence, but also in its expression level [3].

Here we describe the generation of genome-edited alleles of the *bantam* miRNA in *Drosophila*, showing that miRNA seed mutation can profoundly perturb miRNA biogenesis. In that example, a simple cell culture assay can identify miRNA mutants which are inefficiently processed, therefore limiting the risk of preparing mutant animal strains with defective mutant miRNA biogenesis. We applied this method to generate a *Drosophila* strain where the *bantam* seed sequence has been mutated to another hexamer, and where mutant *bantam* accumulates at very similar levels than wild-type *bantam*.

## MATERIALS AND METHODS

Flies were raised under standard culture conditions at 25^*°*^C.

### Molecular biology

Preparative PCR for plasmid construction were performed with Q5 High-Fidelity DNA Polymerase (New England Biolabs cat. #M0491S) following the manufacturer’s protocol. All constructs were verified by Sanger sequencing (Eurofins). Their annotated sequences are available at https://github.com/HKeyHKey/Busseau_et_al_2023/tree/main/Sequences.

### Genome editing

Donor plasmids were prepared by PCR-amplifying chr3L bp 640650–643906 (dm6 assembly) from genomic DNA from *Drosophila* strain nos-Cas9 (Bloomington *Drosophila* Stock Center #54591) with primers d1319 and d1320 (see Supplementary Table 1 for oligo sequences), then cloning that product between the *Sac*I and *Xho*I sites of pBTKS-. Mutant constructs were generated by PCR-mediated mutagenesis on that plasmid, then replacing the *Age*I/*Hin*dIII fragment by the mutated fragment.

sgRNA-expressing plasmids were prepared by cloning oligo duplexes in the *Bbs*I site of plasmid pCFD3-dU6:3gRNA, which was a gift from Simon Bullock (Addgene plasmid #49410; http://n2t.net/addgene:49410; RRID:Addgene_49410; [13]). sgRNA specificity was predicted using CRISPR targetFinder (http://targetfinder.flycrispr.neuro.brown.edu/). For constructs inserted using the PAM located 9 nt downstream of the hairpin, the cloned duplex was made by annealing oligos d1316 and d1343; for constructs inserted using the PAM located 52 nt downstream of the hairpin, the cloned duplex was made by annealing oligos d1938 and d1939.

Edits were performed following the protocols described by [13, 14]: donor plasmid (300 to 700 ng.*µ*L^−1^) and sgRNA producer plasmid (100 to 150 ng.*µ*L^−1^) were co-injected into nos-Cas9 embryos (Bloomington *Drosophila* Stock Center strain BL#54591) using TheBestGene service (https://www.thebestgene.com/) or in our own facility. G0 adults were then crossed to the balancer BL#23232 strain (TM6B-YFP/Dr), their progeny was screened by PCR using oligos specific for the intended mutation, and heterozygous stocks were established and maintained on the TM6B-YFP balancer chromosome from BL#54591. All edited *bantam* alleles were verified by Sanger sequencing (Eurofins).

### Small RNA-Seq

Total RNA was TRIZOL-extracted (Life Technologies) from approximately 100 L1 heterozygous larvae of each genotype, hand-collected under a fluorescent binocular equiped with a GFP filter set. For Figure 1**C**, larvae were collected 0–2 h after hatching; for Figure 4**B**, they were collected 0–6 h after hatching. Samples were then depleted of 2S rRNA by Oligo Capture using biotinylated oligo d2227 and magnetic Dynabeads MyOne Streptavidin C1 (Life Technologies cat. #65001) following the protocol described by [15]. Spike-in RNA oligos were then introduced in the RNA samples: for the experiment shown in Figure 1, an equimolar mix of oligos r25 and r26 was added to each RNA sample (1 *µ*L of a mix containing 5 × 10^−16^ M of each oligo, per *µ*g of total RNA). This spike-in mix was too dilute and spike-in counts were not usable for the four libraries. Another library was then prepared using an aliquot of the “m2(proximal)/+” RNA sample supplemented with a more concentrated oligo mix (1 *µ*L of a mix containing 1.5 × 10^−10^ M of each oligo, per *µ*g of total RNA), and its spike-in counts were used to correct for sequencing biases in the first four libraries.

**Figure 1:**
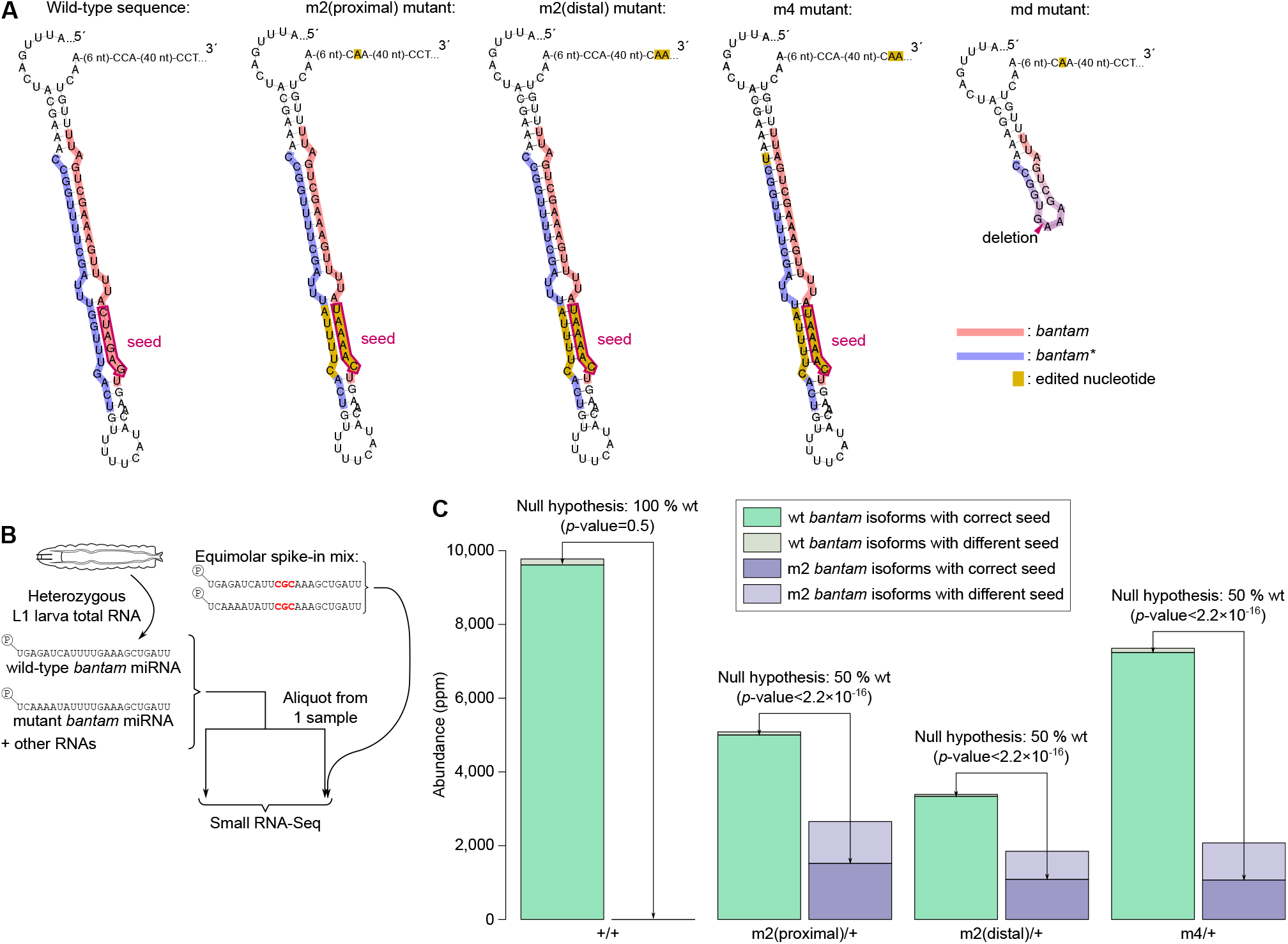
Perturbed miRNA *in vivo* accumulation after seed mutation. **A**. Predicted secondary structures of wild-type and mutated *bantam* loci. Mutated nucleotides are highlighted in yellow. In each construct, mutations were introduced in the miRNA and miRNA* sequences as well as in the PAM used for genome editing, to prevent cleavage of already-edited chromosomes (two distinct PAMs were used, proximal: 9 nt and distal: 52 nt downstream of the *bantam* hairpin). The seed of wild-type *bantam*, and the intended seed of the edited alleles, are framed in dark purple. **B**. Procedure for miRNA quantification. Whole-body L1 larva RNA was analyzed by Small RNA-Seq. An aliquot of one sample (the “m2(proximal)/+” sample) was diverted for sequencing bias measurement: it was supplemented with an equimolar mix of synthetic RNA oligos emulating wild-type and mutant *bantam* miRNAs (except for a trinucleotide tag, here highlighted in red) then analyzed by Small RNA-Seq. Sequencing biases between wild-type and mutant *bantam* were inferred from spike-in read counts in that library, and the observed wild-type/mutant ratio was used to correct miRNA read counts. **C**. Measured abundances of wild-type and m2 *bantam* isoforms in the 4 TM6-balanced L1 larva libraries (the “+/+” control is also heterozygous for the TM6 balancer chromosome). Wild-type isoforms “with correct seeds” are the isoforms having a GAGAUC hexamer at positions 2–7; m2 isoforms “with correct seeds” are the isoforms having a CAAAAU hexamer at positions 2–7. Read counts for m2 *bantam* with correct seed were corrected for sequencing biases using the observed spike-in read counts. “ppm”: parts per million (*i*.*e*.: number of reads per million reads mapping on the genome). Significance of the observed deviations to the expected wt/mutant miRNA ratios was assessed using Fisher’s exact test (null hypothesis and *p*-values indicated for each genotype).

For the experiment shown in Figure 4, an equimolar mix of RNA oligos r25, r26 and r64 was added to each RNA sample (1 *µ*L of a mix containing 1.5 × 10^−10^ M of each oligo, per *µ*g of total RNA).

Small RNA-Seq libraries were prepared by the MGX sequencing facility using the NEXTflex Small RNA-Seq Kit v3 (Bioo scientific), with pre-adenylated 3′ adapter sequence being NNNNTG-GAATTCTCGGGTGCAAGG. Libraries were quantified on a Fragment Analyzer (Agilent) using kit “High sensitivity NGS” and by qPCR on a Roche LightCycler 480.

3′ adapter sequences were trimmed, and reads 18–30 nt long were selected, using cutadapt. Adapter-trimmed reads were mapped by Hisat2 on an index composed of the *Drosophila* dm6 genome, supplemented with sequences of the spiked-in RNA oligos as well as the sequences of the edited *bantam* hairpins. Full-length spike-in sequence occurences were counted in the cutadapt-trimmed fastq files, while wild-type and mutant *bantam* read counts were extracted from the hisat2 mapping output. Read counts were normalized to the total number of small RNA reads mapping on the hisat2 index.

### Luciferase reporter assays

Reporter plasmids containing wild-type, md, m1, m2, m3 or m4 *bantam* variants were constructed by cloning the hairpin of interest (PCR-amplified with oligos d2289 and d2290 from the existing donor DNA, see above) into the *Sac*I and *Xba*I sites of the pmirGLO vector (Promega cat. #E1330).

Reporter plasmids containing m5, m6, m7, m8 or m9 variants were constructed using NEBuilder HiFi DNA Assembly (NEB cat. #E2621) by assembling a PCR fragment amplified using oligos d2436 and d2437 from the reporter plasmid pmirGLO containing wild-type *bantam*, together with oligo duplexes made by annealing oligos d2438 and d2439 (m5), d2440 and d2441 (m6), d2442 and d2443 (m7), d2444 and d2445 (m8), or d2446 and d2447 (m9).

S2R+ cells [16] from DGRC were cultivated under standard conditions at 25^*°*^C in Schneider medium with 10% Foetal Bovine Serum and 1% penicillin-streptomycin. Cells were seeded at 3 × 10^5^ cells.well^−1^ in 6-well plates and transfected 24 h later with one pmiRGLO derivative construct per well (0.9 *µ*g plasmid) using Effectene (Qiagen cat. #301425). After 3 days the cells were washed with PBS and lysed with Cell Culture Lysis Buffer (Promega cat. #E1531) prior to firefly and Renilla luciferase activity measurements which were performed in triplicate using the Dual-GLO kit (Promega cat. #E2920). The whole procedure was repeated three times independently for each construct.

### Gene knockdown

RNA interference experiments to knock down Drosha and Exportin-5 (also known as RanBP21) in S2R+ cells were performed as described by [17] with the following modifications.

Gene fragments for the preparation of dsRNA were obtained using genomic DNA of the transgenic fly strain containing the UASp-GFP-bantam reporter as a template to generate PCR products with flanking T7 RNA polymerase promoters using oligonucleotides d2016 and d2017 (685 nt of Drosha sequence), d2018 and d2019 (718 nt of Exportin-5 sequence), d2630 and d2631 (666 nt of GFP sequence).

RNA synthesis from the PCR products was performed using the Megascript T7 kit (Life Technologies) following exactly the manufacturer’s procedure including DNAse I treatment followed by LiCl precipitation. The quality of dsRNA was estimated by standard agarose gel electrophoresis and the concentration was determined on a Nanodrop spectrophotometer.

S2R+ cells were seeded at 10^6^ cells.mL^−1^ in 5 mL medium in a T25 flask, and 50 *µ*g dsRNA was added directly to the growth medium. After four days, an additional 50 *µ*g dsRNA was added, and the cells were diluted 2-fold into 10 wells of a 12-well plate (1 mL medium per well). After two days they were transfected and an additional 10 *µ*g dsRNA was added per well. Transfections were performed in duplicate into 8 of the wells with pmiR-GLO-banmd, banwt, banm2 and banm6, and two wells were mock-transfected. Eight days after the initial dsRNA treatment, cells transfected with pmiR-GLO derivatives were processed for Luciferase assays, and the mock-transfected cells were washed with PBS then scrapped in 1 mL TRIZOL (Thermo Fisher Scientific) for RNA extraction, followed by DNase I treatment (RQ1, Promega) and ethanol precipitation. The quality of RNA was estimated by standard agarose gel electrophoresis and the concentration was determined on a Nanodrop spectrophotometer.

One microgram total RNA was reverse transcribed using oligo-dT 12–18 nt long (Life Technologies) and SuperScript II reverse transcriptase (Life Technologies). Quantitative PCR was performed using the LightCycler 480 SYBR Green I Master system (Roche) with the housekeeping genes Actin 42A and Tubulin 84B as references. Primers were d2638 and d2639 (Drosha), d2642 and d2637 (Exportin-5), d1654 and d1655 (Actin 42A), and d1658 and d1659 (Tubulin 84B).

### *In vivo* GFP reporter assays

The GFP reporter plasmid containing a wild-type *bantam* hairpin in its 3′ UTR was constructed as follows: a GFP-*bantam* fragment was amplified using oligos d1360 and d1361 from the transgenic fly strain containing UAS-banD [18], digested with *Bam*H1 and *Xba*1, and ligated together with a *Xba*I-*Nde*I fragment containing the SV40 3′ UTR from vector pUASt-attB [19] into *Bam*H1-*Nde*I-digested vector pUASpK10 [20]. GFP reporter plasmids containing md, m2 and m6 were generated by replacing the *Mlu*1-*Xba*1 fragment containing the *bantam* hairpin by a *Mlu*I-*Xba*I-digested PCR fragment generated using oligos d1379 and d1380 from the corresponding existing donor plasmids.

PhiC31 integrase-mediated transgenesis was performed by the BestGene facility. Transgenes were targetted into the attP2 docking site of stock BL#8622.

Tissue-specific expression of the GFP-reporter transgenes in the female germline was induced using the nos-Gal4-VP16 driver [21]. Ovaries were hand-dissected from 3 day-old females maintained on medium supplemented with dry yeast, fixed in 4% PFA and stained with 1 *µ*g.mL^−1^ DAPI following standard procedures, then mounted in Vectashield. Images were captured with a Zeiss Axioimager Apotome microscope.

### Data and script availability

Small RNA-Seq data has been deposited at NCBI’s SRA (https://www.ncbi.nlm.nih.gov/sra/) under accession numbers SUB8281052 (for Figure 1) and SUB11510745 (for Figure 4). Scripts and intermediary files are accessible at https://github.com/HKeyHKey/Busseau_et_al_2023.

## RESULTS

In an effort to generate a series of mutated alleles of the *Drosophila bantam* miRNA gene, we used CRISPR/Cas9-mediated genome editing to prepare three *Drosophila* lines where the *bantam* seed is mutated, and a fourth line where the *bantam* pre-miRNA hairpin is truncated (see Figure 1**A**). In each seed mutant, miRNA* nucleotides facing the miRNA seed nucleotides were mutated too in order to preserve pre-miRNA secondary structure.

miRNA expression was then assessed by Small RNA-Seq in L1 larvae, where *bantam* is expressed at high levels [18]. We analyzed larvae heterozygous for the wild-type allele to provide an internal control. Biases in the propensity of various small RNAs to be deep-sequenced may distort their abundances in the Small RNA-Seq libraries [22, 23]. In order to measure and correct such biases between wild-type and mutant *bantam*, RNA samples were mixed with spiked-in synthetic RNA oligos exhibiting the same sequence as the wild-type and intended mutant miRNA except for a central trinucleotide tag distinguishing them from miRNAs expressed from the larvae (see Figure 1**B**).

The replacement of the *bantam* seed sequence (as well as facing *bantam*\* sequences, and the PAM located 9 nt downstream of the stem-loop) greatly reduced miRNA accumulation (compare wt to m2 *bantam* abundances for m2(proximal)/+ heterozygotes on Figure 1**C**: 70% reduction for the mutant allele). It also perturbs the balance between miRNA isoforms: while wild-type *bantam* expresses mostly isoforms sharing the same 5′ extremity, hence the same seed sequence, m2 *bantam* is expressed as a balanced mixture of isoforms with the intended CAAAAU seed and isoforms with alternative seed sequences (compare dark and pale purple on Figure 1**C**). Because it is conceivable that the PAM mutation disrupts some unknown important sequence or structure motif, we prepared another version of the m2 edited hairpin, keeping that PAM unchanged and mutating another, distant PAM. That modification did not improve mutant miRNA accumulation (see m2(distal)/+ on Figure 1**C**: 67% reduction for the mutant allele). In an attempt to reinforce miRNA/miRNA* asymmetry, we introduced an additional mutation, strengthening duplex base-pairing at the miRNA* 5′ end [24, 25, 26]. While that mutation was predicted to increase miRNA accumulation, here it proved ineffective since m4 heterozygotes do not accumulate more mutant miRNA than m2 (see m4/+ on Figure 1**C**: 85% reduction for the mutant allele).

We reasoned that the deleterious effect of the seed mutation may derive from a destabilization of the pre-miRNA stem double-helix: while the wild-type *bantam* seed contains 3 (C+G) and 3 (A+U) nucleotides, the mutant seed contains 1 (C+G) and 5 (A+U) nucleotides. In mammals, Drosha-mediated cleavage requires the double helix to be long enough, at least because its partner DGCR8 needs to bind to a stem of sufficient length [10]. It is therefore possible that a (A+U)-rich *bantam* seed decreases the pairing stability of the stem to the point that it is less efficiently recognized by the Drosha machinery.

We therefore aimed at designing a test to monitor the ability of candidate hairpins to be cleaved by Drosha in cultured cells, similar to the assay described by [27]. The insertion of the wild-type *bantam* hairpin in the 3′ UTR of a firefly luciferase reporter inhibits reporter expression in S2R+ cells, relatively to the insertion of the truncated “md” hairpin, which is unsuitable for Drosha processing (see Figures 2**A** and **B**). These results are in agreement with the notion that Drosha-mediated cleavage promotes degradation of the host transcript (reviewed in [28]), and provides the basis for a simple *ex vivo* assay to estimate the ability of candidate pri-miRNAs to serve as Drosha substrates.

**Figure 2:**
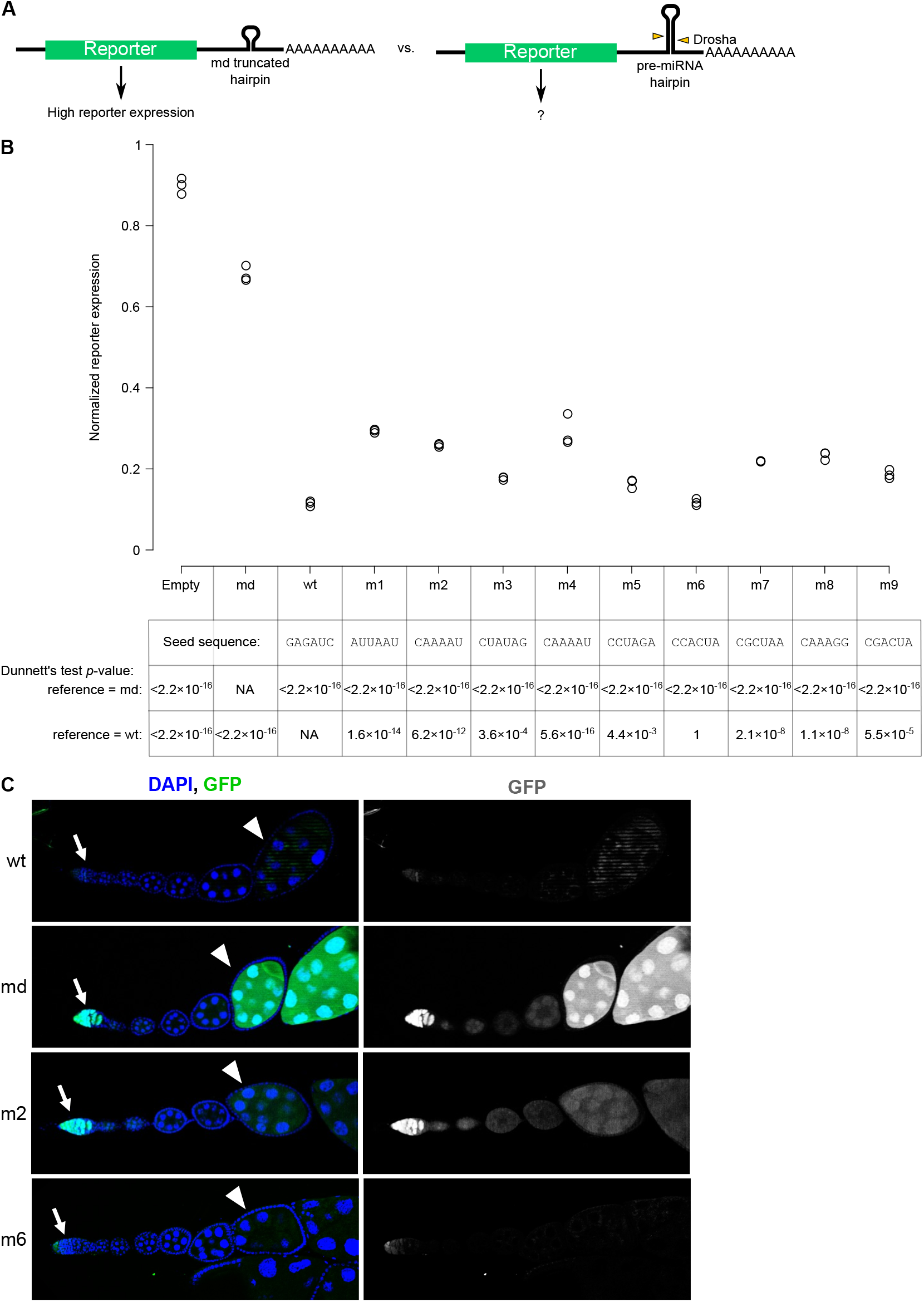
*Ex vivo* screen for an estimation of Drosha substrate suitability. **A**. Principle of the assay: insertion of a candidate pre-miRNA hairpin in the reporter’s 3′ UTR attracts Drosha cleavage more or less efficiently, resulting in various levels of reporter expression. Efficiently processed hairpins induce more degradation of the host transcript, hence a lower reporter expression. **B**. Candidate hairpins were inserted in the 3′ UTR of a firefly luciferase reporter, expressed from the same plasmid than a Renilla luciferase internal control. For each construct, the sequence of its mutated seed is given in the table under the graph. After subtraction of luciferase signals measured in untransfected cells, the firefly/Renilla luciferase activity ratio reveals the repressive action of inserted hairpins on firefly expression. The experiment was performed in three biological replicates, and firefly/Renilla luciferase ratios were compared to the ratio measured in two reference transfections (with an inserted “md” truncated hairpin, and with a wild-type *bantam* hairpin); Dunnett’s test *p*-values are indicated in the table under the graph. **C**. Candidate hairpins corresponding to *bantam* wt, md, m2 and m6 were inserted in the 3′ UTR of a a UASp-GFP reporter. Expression was driven in the female germline using nos-GAL4-VP16, whose two peaks of activity reside in the germ cells of the germarium (arrows) and the developing stage 6–7 egg chambers (arrowheads).

We therefore compared 9 distinct mutant pri-miRNA sequences for their ability to repress a hostluciferase reporter gene. Sequence and predicted secondary structure for these 9 variants are shown in Supplementary Figure 1. We detected a significant reduction of luciferase expression for each of these candidates compared to the md hairpin, indicating that they all undergo some level of cleavage by Drosha (see Figure 2**B**). When compared to the wild-type hairpin, every candidate was significantly less efficient at repressing their host reporter gene, except for the m6 construct (see Figure 2**B**), suggesting that the m6 allele may be matured with an efficiency similar to that of wild-type *bantam*.

Consistent with the notion that inserted pre-miRNA hairpins inhibit reporter expression because of Drosha-mediated degradation of the reporter mRNA, we observed that this effect is essentially abolished when cells have been depleted for Drosha by RNAi (see Figure 3). In a control experiment, we also depleted Exportin-5 by RNAi before transfecting the reporter plasmid. Exportin-5 (also known as Ranbp21 in *Drosophila*) is required for nucleo-cytoplasmic export of pre-miRNAs in vertebrates [29, 30, 31], hence downstream of Drosha-mediated cleavage of the pri-miRNA. Surprisingly still, its knockdown also interfered with the inhibition of reporter expression by inserted hairpins (see Supplementary Figure 2). This unexpected effect may be due to the role of *Drosophila* Exportin-5 in tRNA export, which could impact firefly and Renilla luciferase translation [32].

**Figure 3:**
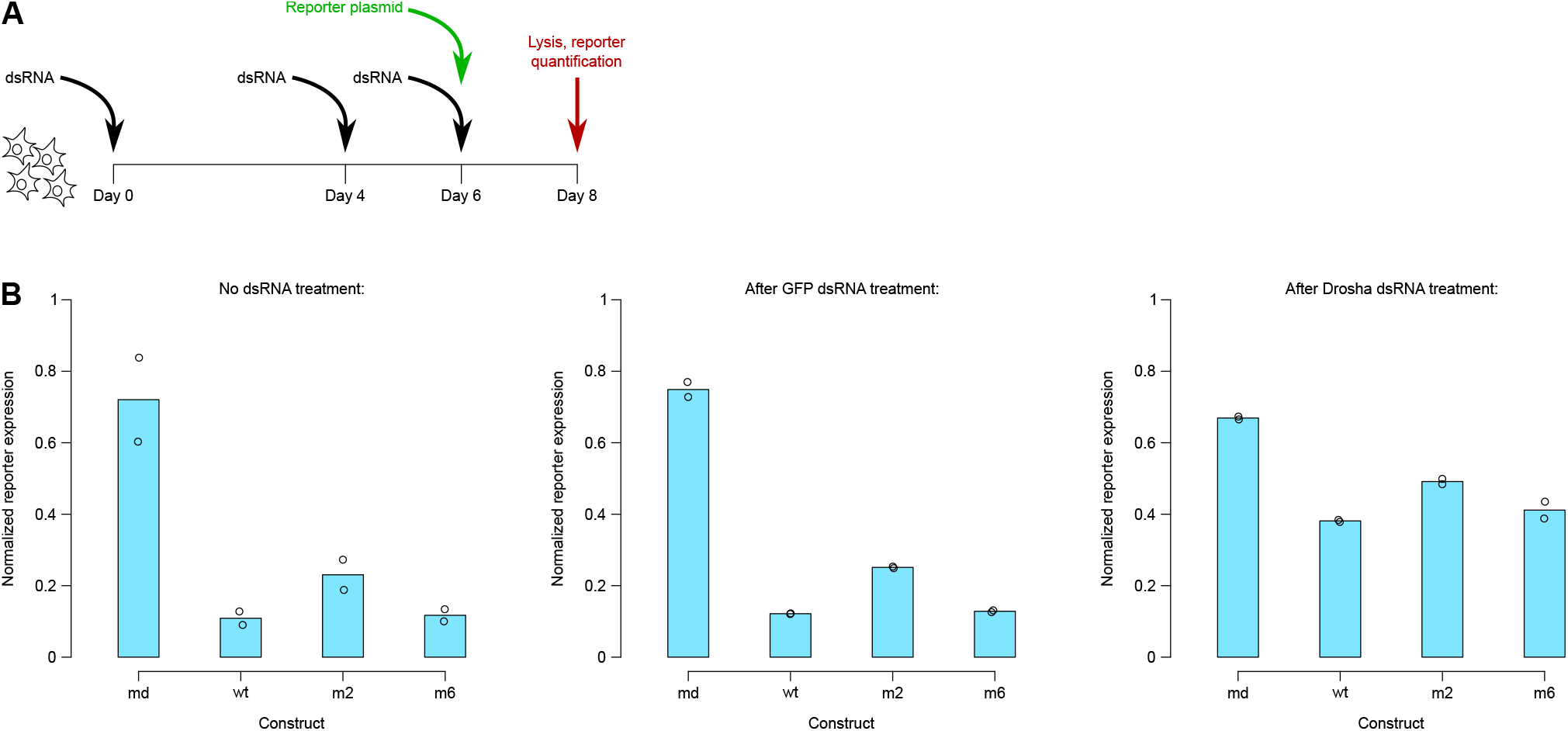
Hairpin insertion-mediated inhibition of reporter expression is Drosha-dependent. **A**. Experimental procedure. Target genes were repressed by dsRNA-mediated RNAi, with repeated addition of dsRNA in the cell medium. The reporter plasmid (expressing firefly luciferase with various hairpins inserted in its 3′ UTR, as well as a normalizing Renilla luciferase) was transfected 2 days before cell collection and reporter quantification. **B**. Firefly/Renilla luciferase activity ratio (*y* axis) depends on the identity of the hairpin inserted in firely luciferase 3′ UTR (md, wt, m2 or m6; on the *x* axis), and this effect is mostly lost when Drosha had been repressed by RNAi. An ANOVA analysis shows that hairpin identity (*p*=3. × 6 10^−9^), dsRNA identity (*p*=1.6 × 10^−5^) as well as the interaction between these two factors (*p*=0.0030) have a significant effect. Two biological replicates of the reporter assay were prepared from a common culture of dsRNA-treated cells.

**Figure 4:**
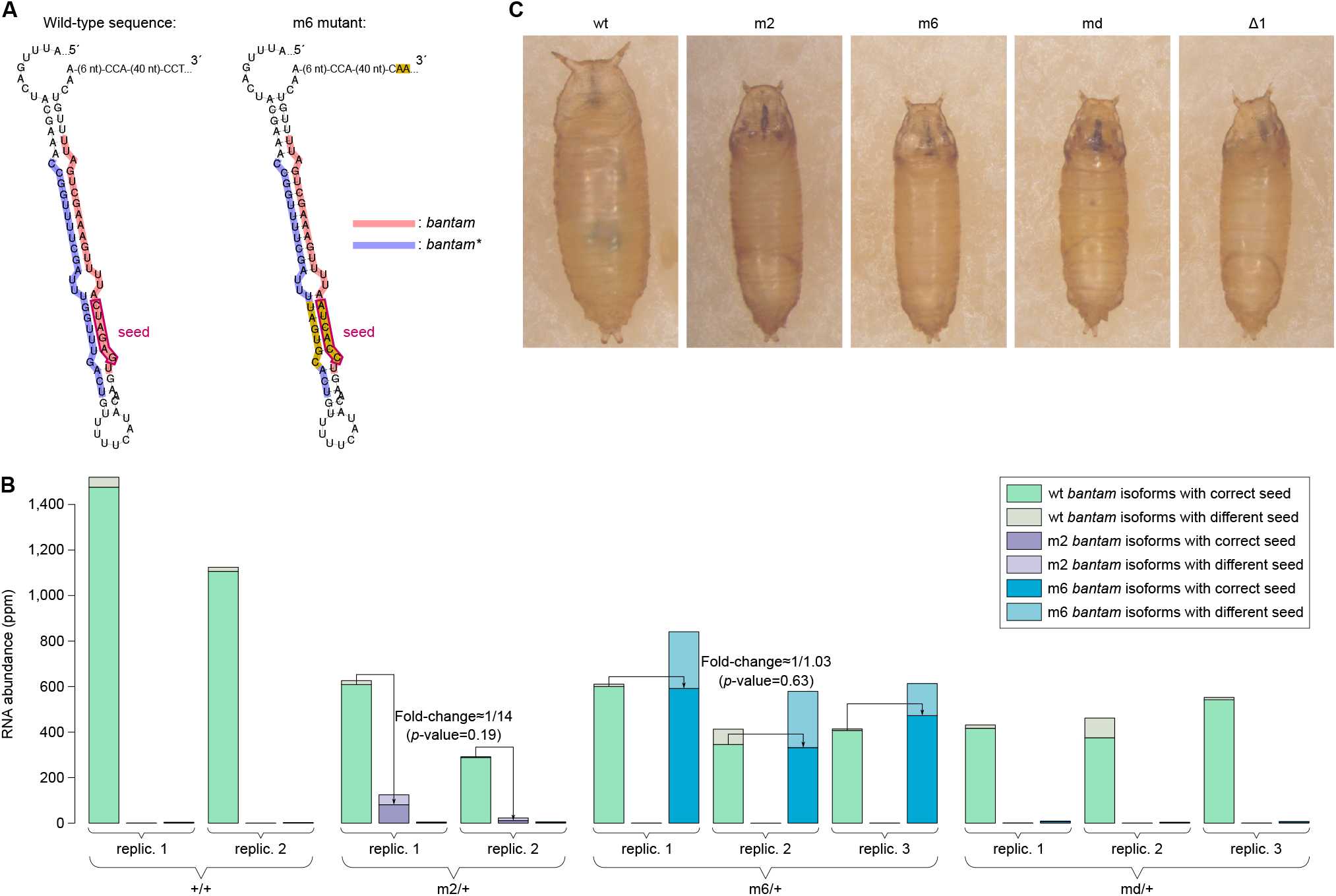
Generation of an edited *bantam* allele with correct expression level. **A**. Predicted secondary structure of wild-type and “m6” edited *bantam* loci. Same conventions as in Figure 1**A. B**. Measured abundances of wild-type, m2 and m6 *bantam* isoforms in TM6-balanced heterozygous L1 larvae (the “+/+” control is also heterozygous for the TM6 balancer chromosome). Wild-type isoforms “with correct seeds” are the isoforms having a GAGAUC hexamer at positions 2–7; m2 isoforms “with correct seeds” are the isoforms having a CAAAAU hexamer at positions 2–7; m6 isoforms “with correct seeds” are the isoforms having a CCACUA hexamer at positions 2–7. Read counts for m2 and m6 *bantam* with correct seed were corrected for sequencing biases using the observed spike-in read counts. Measured fold-changes between wild-type and m2 or m6 alleles are indicated (geometric mean of the fold-change across 2 or 3 biological replicates for m2/TM6 and m6/TM6 genotypes respectively; *p*-values were calculated using a paired t-test). **C**. Growth defects in wild-type, homozygous *bantam*-edited strains, and in Δ1 homozygous pupae [33], imaged at the same magnification.

We also inserted *bantam* hairpin variants in a GFP host gene to assess their repressive effect *in vivo* in *Drosophila* ovaries (see Figure 2**C**). The reporter with the uncleavable md variant produced high GFP both in the germarium and developing egg chambers, while the reporter with the m2 variant produced lower level of GFP in egg chambers (see Figure 2**C**). In contrast, GFP from the reporters with wt *bantam* or the m6 variant was barely detectable both in the germarium and in developing egg chambers. These *in vivo* results mirror the *ex vivo* data, confirming that the presence of a m6 or wt hairpin in the 3′ UTR of a reporter mRNA decreases its expression to a similar extent, while the m2 variant exerts a weaker repressive effect.

These results prompted us to prepare a *Drosophila* strain where the *bantam* gene has been mutated into the m6 variant (see Figure 4**A**). Whole-body RNA from heterozygous L1 larvae was analyzed by Small RNA-Seq, using spiked-in RNA oligos for bias correction in each library. In order to take into account any potential difference in miRNA content across independent batches of≈ 100 larvae, in this experiment we analyzed several replicates of larva batch. This design allowed us to not only compare the relative abundance of mutant and wild-type miRNA alleles within heterozygotes, but also to estimate the reproducibility of the observed fold-changes across independent batches of larvae. The availability of 10 spiked-in libraries also allowed us to verify the reproducibility of sequencing biases between the wild-type, m2 and m6 alleles of the *bantam* miRNA: ratios between the raw read counts for wt, m2 and m6 oligos proved consistent from library to library (see Supplementary Figure 3).

In terms of abundance of miRNA isoforms with the correct seed sequence, expression of the m6 allele is very similar to that of the wild-type allele, and we were unable to detect a significant difference between the abundances of these two miRNA sequences (see Figure 4**B**). The m6 allele therefore constitutes a successfully-edited allele, where the *bantam* seed has been mutated into a different hexamer without noticeably perturbing its accumulation level. The m2 allele, which was re-analyzed in parallel, shows a greater difference with the wild-type allele, although the observed fold-changes were quite variable and did not prove reproducibly different from 1 – likely because we could only analyze two replicates in that genotype.

Note that *bantam* abundance differs between Figure 4**B** and Figure 1**C** because, for practical reasons due to the number of replicates to be collected, L1 larvae were not at the exact same stage between the two experiments (larvae analyzed in Figure 1**C** tended to be younger than those analyzed in Figure 4**B**, and *bantam* expression decreases very fast during early larval development: [18]).

The availability of this allelic series allowed us to evaluate the phenotypes of homozygous edited *bantam* mutants. Both the m2 and m6 seed mutants, as well as the md deletion mutant, recapitulated the reported *bantam* null (called “Δ1”) phenotypes [33, 18]: larval growth defects (see Figure 4**C**), and larval and pupal lethality (not shown). This observation confirms that mutating the miRNA seed is sufficient to phenocopy a miRNA null mutant, regardless of mutant miRNA expression level.

## DISCUSSION

Genome editing holds great promise for the functional assessment of individual miRNA/target interactions. Allowing the mutation of miRNA genes, or of their binding sites in target RNAs, or both of them simultaneously in a compensatory manner, this technique should allow miRNA/target interactions to be disrupted and restored on demand in a complete *in vivo* setting. The analysis of physiological phenotypes controlled by individual interactions is much needed, as exemplified by the frequent mis-annotation of the biological roles of miRNA/target interactions when it is based on extrapolated *ex vivo* or *in silico* analyses [34, 35].

Yet the interpretation of physiological phenotypes in such miRNA edited mutants is complicated by the influence of miRNA sequence on its accumulation level [3]. It is therefore essential to generate miRNA mutant strains where not only the miRNA sequence has been edited, but its accumulation level is as close as possible to the wild-type miRNA abundance. The cell culture screening method that we describe here allowed us to identify a mutant construct which emulates wild-type miRNA expression levels for a seed-edited miRNA. Our method is easily generalizable to other miRNAs and other species.

This assay approximates Drosha cleavage efficiency by the reduction of expression of a host reporter gene. Additional mechanisms may alter reporter expression under the control of the inserted pre-miRNA hairpin sequences, therefore potentially blurring the prediction of Drosha cleavage efficiency. Also, miRNA expression levels depend on additional factors (transcription, Dicer cleavage, loading on the effector protein, post-loading stability, …), which can also be influenced by miRNA sequence, and which are not captured by our assay. In particular, accumulated miRNA/miRNA* ratios appear to differ between wild-type, m2 and m6 alleles (see Supplementary Figure 4: miRNA/miRNA* ratio is ≈ 15 for the wild-type allele, and≈ 3 and *≈* 6 for m2 and m6 alleles respectively). This observation shows that the efficiency of Drosha-mediated maturation is one factor among others, controlling the accumulation of seed-edited miRNAs.

We also observed that the mutation of the miRNA seed sequence (together with that of facing miRNA* nucleotides) affects the distribution of miRNA isoforms. While wild-type *bantam* harbors a very homogeneous 5′ end (yielding mature molecules which have mostly the same seed sequence), we note that m2 and m6 mutants tend to generate more heterogeneous 5′ ends (see Figures 1**C** and 4**B**). This result reveals a subtle dependence of Dicer cleavage site selection, of miRNA loading efficiency, or of miRNA stability, to miRNA sequence, even without any change in the predicted hairpin secondary structure. In the case of the *bantam* m6 mutant, a simultaneous increase in pan-isoform miRNA accumulation compensates for this effect, therefore achieving a highly accurate wild-type level of accumulation for the m6 isoforms with correct seed sequence. Further analyses will be required to characterize more finely such subtle consequences of sequence editing on various aspects of miRNA biogenesis and accumulation.

## CONCLUSIONS

Editing *bantam*’s seed sequence triggered unwanted perturbation of miRNA biogenesis *in vivo*, even though facing miRNA* nucleotides were simultaneously mutated in an attempt to preserve pre-miRNA secondary structure. Screening candidate edited pre-miRNA sequences in a cultured cell assay helped identifying the best processed candidate, therefore allowing us to generate an edited *Drosophila* line where the *bantam* seed has been mutated without affecting its accumulation level. Usage of this assay could be generalized, to screen candidate mutant pre-miRNAs for their ability to be efficiently processed.

## ACKNOWLEDGMENTS

We are grateful to Simon Nicot for guidance in cell culture, Lara Demont for help in the screening of edited flies, Laly Robledo for her implication in the setting of Luciferase assays, and Guillaume Canal for his help in optimizing spike-in oligo sequences. We acknowledge the MRI imaging facility, member of the France-BioImaging infrastructure, and the MGX sequencing facility, member of the France Génomique National Infrastructure, supported respectively by contracts ANR-10-INBS-04 and ANR-10-INBS-09 “Investments for the future” of French National Research Agency, as well as the Biocampus *Drosophila* facility. Stocks obtained from the Bloomington Drosophila Stock Center (NIH P40OD018537) were used in this study. This work was funded by the CNRS.

## AUTHOR CONTRIBUTION

B. and H. Seitz conceived the project; I. B., É. H. and H. Somaï collected data; S. M. and H. Seitz analysed data; I. B. and H. Seitz wrote the manuscript. All authors have approved the final article.

## SUPPLEMENTARY DATA

**Supplementary Figure 1:**
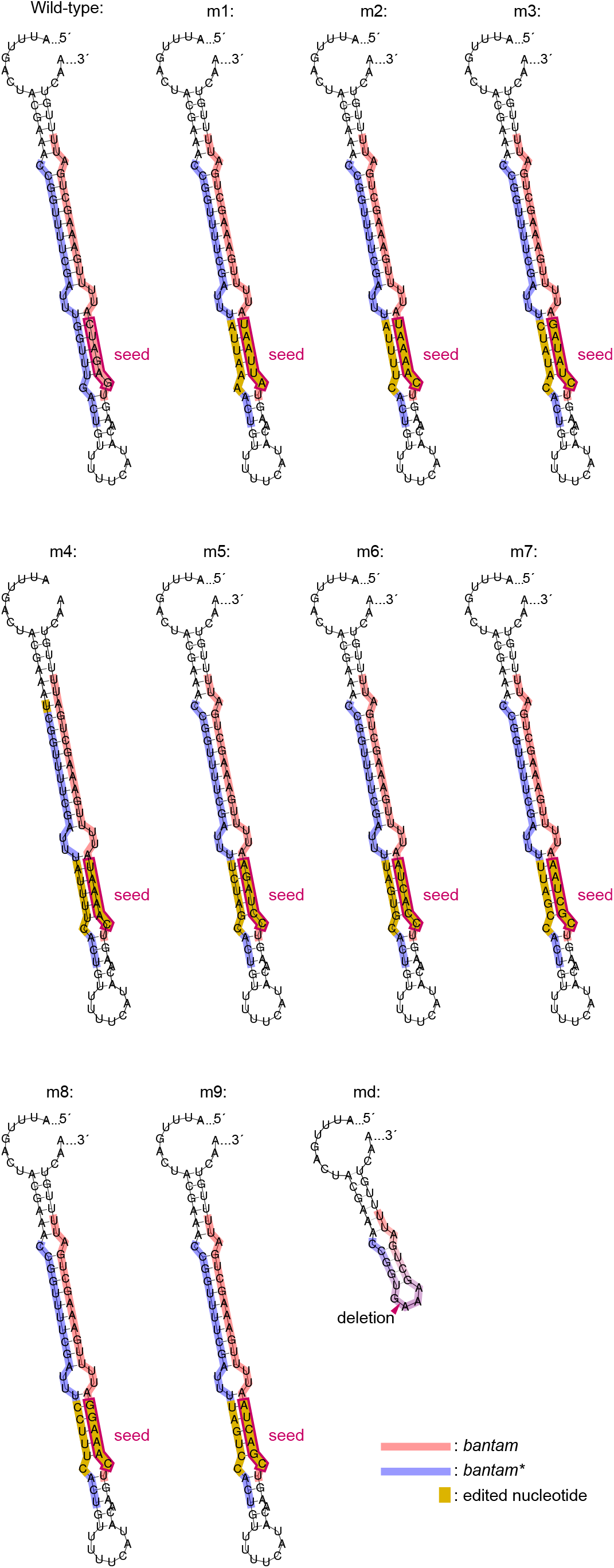
Sequence and predicted secondary structure for the variants analyzed in Figure 2. Same conventions as in **Figure 1A**.

**Supplementary Figure 2:**
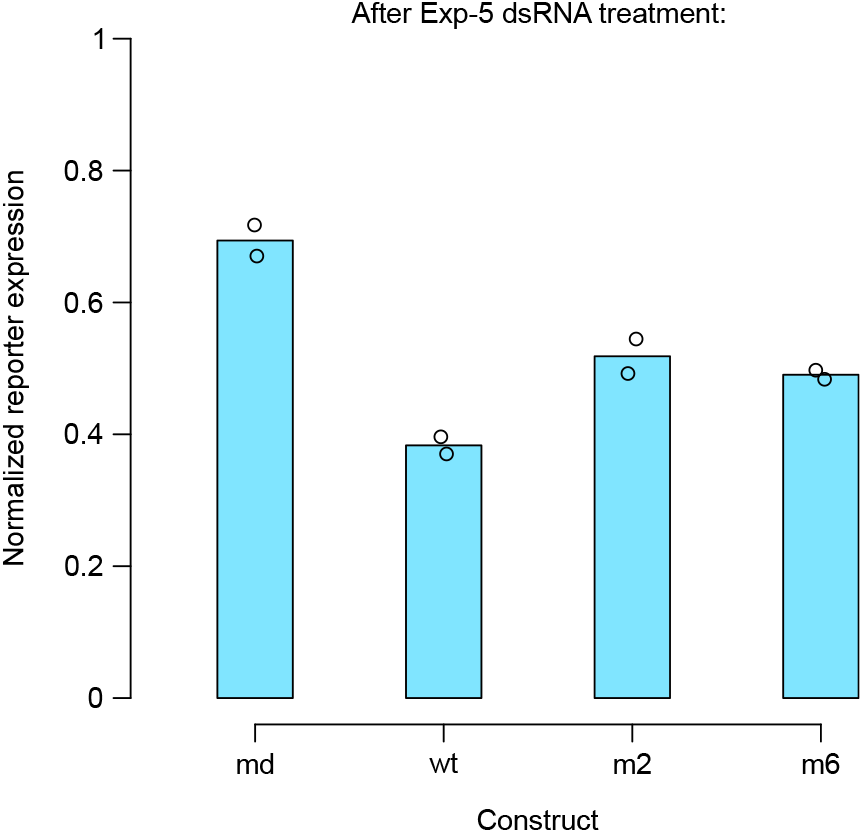
Exportin-5 knockdown interferes with hairpin insertion-mediated inhibition of reporter expression. Same conventions as in Figure 3.

**Supplementary Figure 3:**
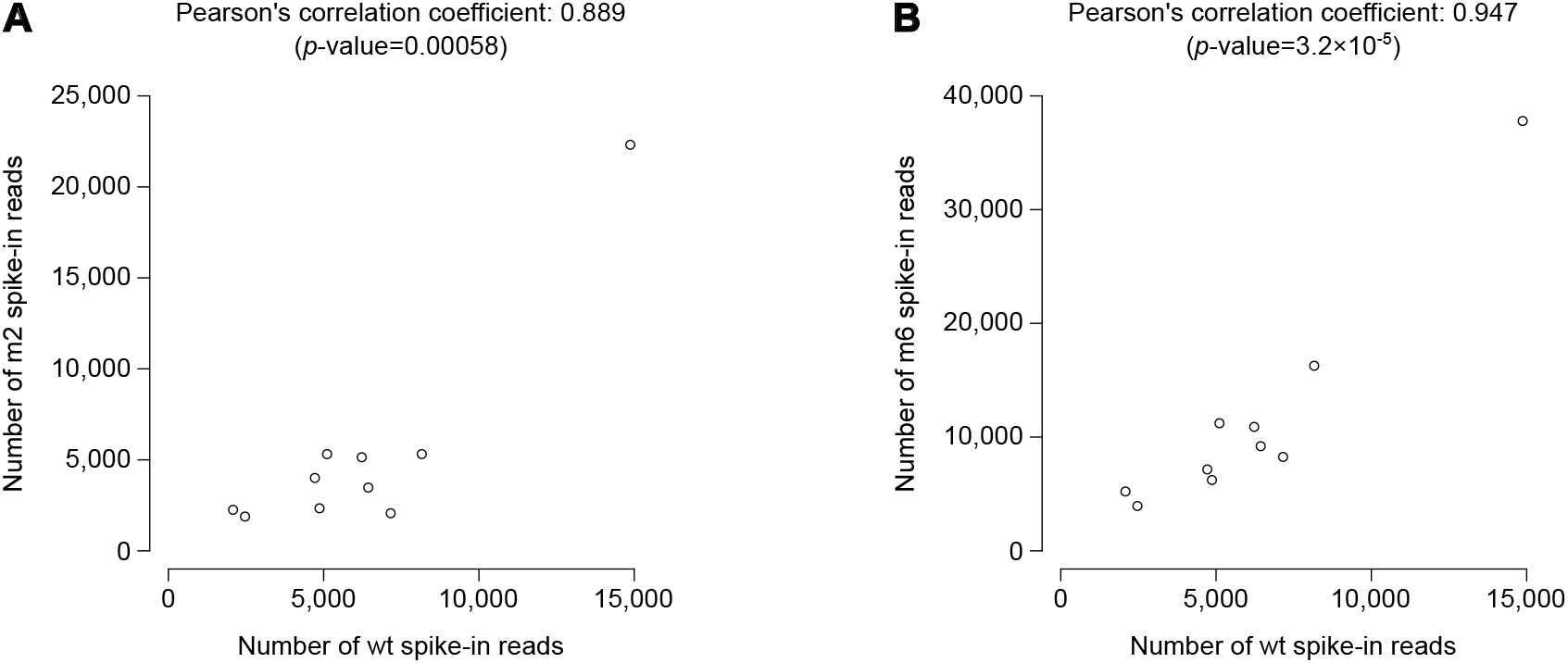
Reproducibility of sequencing biases among full-length wild-type, m2 and m6 *bantam* miRNAs. Raw read counts for spike-ins were measured in the 10 libraries shown in **Figure 4B** (each point represents one library). **A**. *x*-axis: read count for the wild-type *bantam* spike-in; *y*-axis: read count for the m2 *bantam* spike-in. **B**. *x*-axis: read count for the wild-type *bantam* spike-in; *y*-axis: read count for the m6 *bantam* spike-in.

**Supplementary Figure 4:**
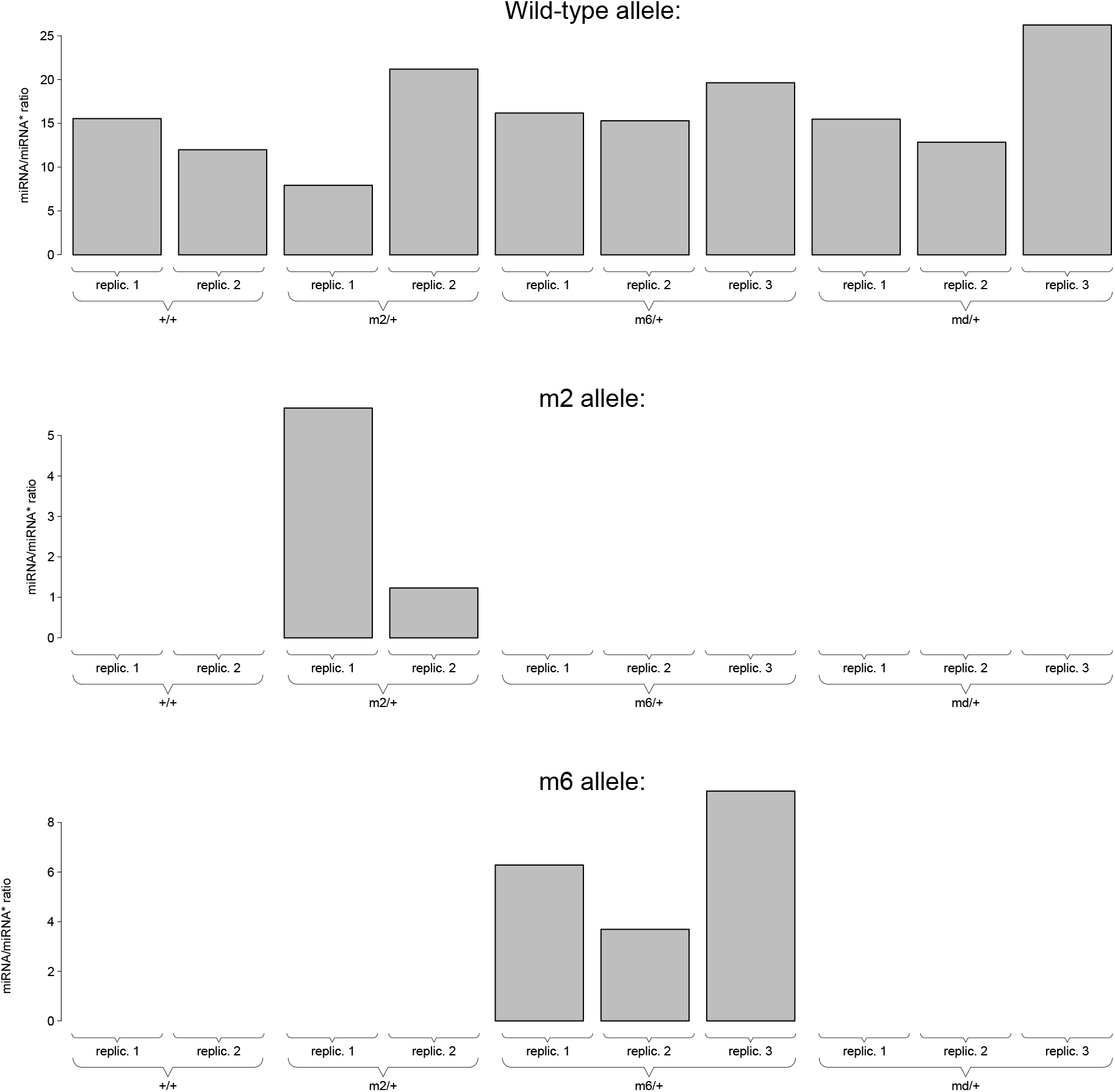
miRNA/miRNA* abundance ratios for wild-type, m2 and m6 *bantam* alleles. Raw read counts for *bantam* miRNA and *bantam\** miRNA* were measured in every library shown in Figure 4, and displayed for every relevant library (*i*.*e*.: all 10 libraries are shown in the “wild-type allele” panel at the top; only m2/+ and m6/+ libraries are shown in the middle and bottom panels respectively). ANOVA *p*-value for the effect of allele identity on miRNA/miRNA* ratio: 0.0034.

**Supplementary Table 1:**
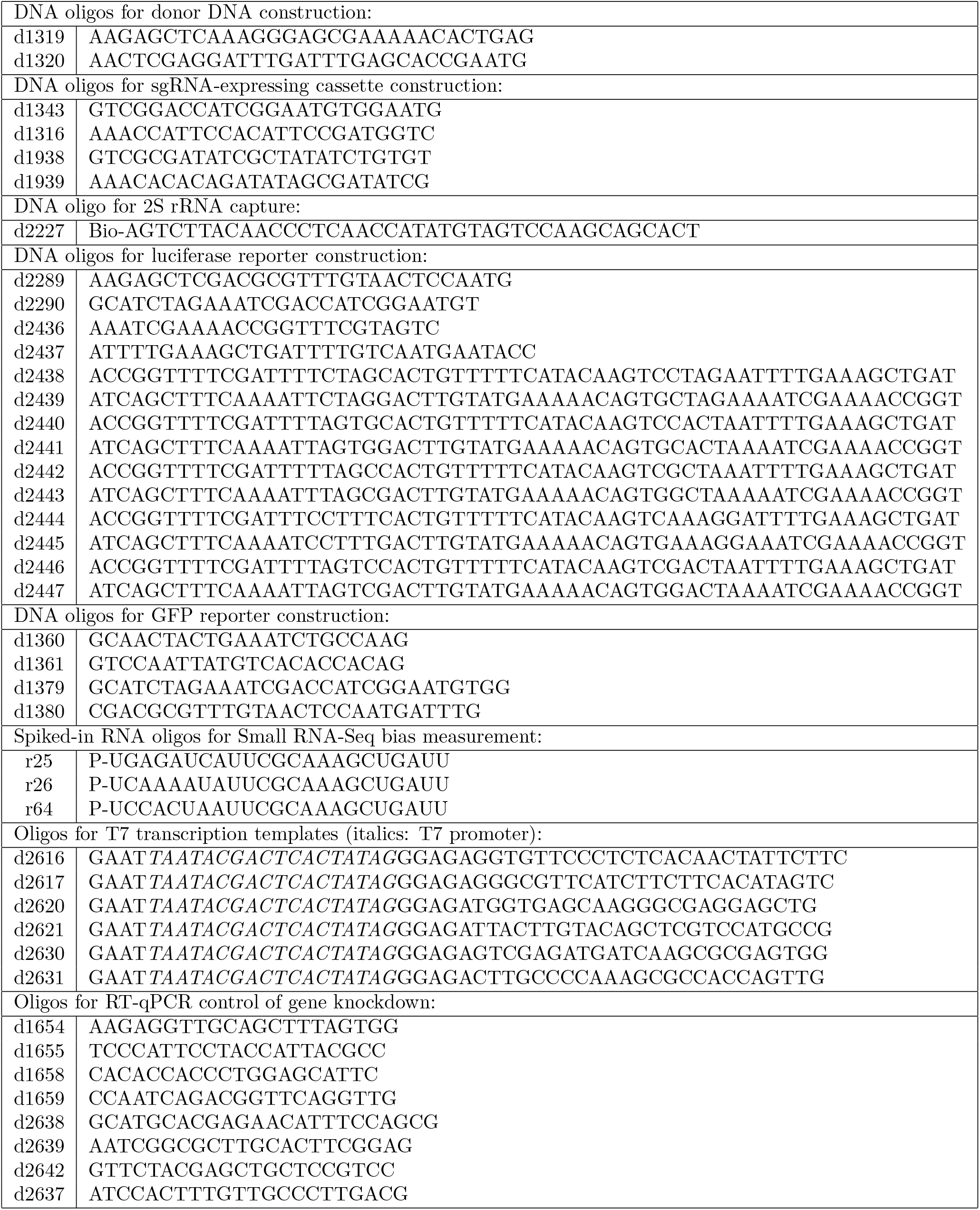
Oligonucleotides used in this study. “Bio”: biotin (with a 11-heteroatom linker to 5′ phosphate). “P”: phosphate. Sequences are written 5′ to 3′.

## Notes

### Competing Interest Statement

The authors have declared no competing interest.

### Summary of Updates

Added statistical analysis for Fig. 1C. Clarifying a few points (aim of spike-in-based correction, comparison of phenotypes upon seed edition and null mutation, evaluating imbalance between miRNA and miRNA* in edited and wt alleles, verification of Drosha-dependency of the observed differences between alleles). Correcting a few grammar issues. Adding "Conclusions" and "Author contribution" sections.

